# Chemotaxonomy of *Tapinoma* and some Dolichoderinae ants from Europe and North Africa

**DOI:** 10.1101/2022.09.28.509850

**Authors:** Alain Lenoir, Elfie Perdereau, Laurence Berville

**Affiliations:** IRBI Institut de Recherche sur la Biologie de l’Insecte, UMR CNRS 7261, Université de Tours, Faculté des Sciences, Parc de Grandmont, 37200 Tours, France

**Keywords:** Chemosystematics, Cuticular hydrocarbons, invasive ants, *Tapinoma*, *Dolichoderus*, *Linepithema*, *Bothriomyrmex*

## Abstract

Cuticular hydrocarbons of some Dolichoderinae species from France and various places like Spain, North Africa, and Italy were studied. The *Tapinoma nigerrimum* group was particularly analyzed and replaced in the genus *Tapinoma*. All species were correctly discriminated and a new hydrocarbons profile was found in Spanish mountains in the *T. nigerrimum* group, which was provisionally named *T. sp. Spain*. We added numerous unknown spots for the distribution of these ants. We also tested aggression between some *T. magnum* colonies and it appeared that this species forms supercolonies like other invasive species, but does not form giant supercolonies like the Argentine ant

## Introduction

Cuticular hydrocarbons profiles (CHs) are a good indicator of species discrimination in insects. In ants, Martin and Drijfhout (2009) found more than 1 000 hydrocarbons in 78 ant species and each species possess its own unique pattern. In 12 species of European *Myrmica* Guillem (et al 2016) found remarkable species-specific chemical profiles. On 2 *Temnothorax* and 2 *Myrmica* species, Sprenger and Menzel (2020) assigned the right species based on HCs with 0% errors. In some cases, cryptic species could be discriminated for example in *Tetramorium* (Cordonnier et al 2018), and in tropical arborical parabiotic species (Hartke et al 2019). Peña-Carrillo et al (2021) also found different cryptic species in *Ectatomma ruidum* (Roger, 1860). In some species, colonies from different localities can have different profiles which may indicate different species. For instance, Dahbi et al (1996) found distinct HCs profiles for *Cataglyphis iberica* (Emery, 1906) between Barcelona and Murcia, and the population from Murcia was later described as a distinct species, called *Cataglyphis gadeai* (De Haro and Collingwood, 2003). It has been confirmed later with molecular biology (Villalta et al 2018). On the contrary, some species like *Lasius niger* do not change hydrocarbon profile according to all their European distribution (Lenoir et al 2009).

Dolichoderinae is a large subfamily of ants with approximately 900 described species (Ward et al 2010). They are commonly referred to as odorous ants, referencing to the volatile compounds reminiscent of fermented cheese or rotting fruit emitted from their pygydial (anal) gland (Penick and Smith 2015 for *Tapinoma sessile* (Say, 1836)). In France it is called rancid butter odor.

The taxonomy of the genus *Tapinoma* has been recently reviewed (Seifert 2012), and more recently the *T. nigerrimum* group was separated into four cryptic species (*T. darioi* Seifert et al., 2017, close to *T. magnum* Mayr, 1861, *T. ibericum* Santschi, 1925, *T. nigerrimum* (Nylander, 1856) *sensu stricto* (Seifert et al 2017). A chemical analysis of glandular volatiles confirmed the separation between *T. darioi* and *T. magnum* (D’Eustachio et al 2019). Nevertheless, some subspecies can now be separated in two good species using DNA as for example *Tapinoma atriceps* and *T. atriceps breviscapum* in Brazil (Escárraga et al 2021).

Cuticular hydrocarbons of *Tapinoma* have been investigated previously only in a few species: *T. erraticum* (Latreille, 1798), *T. israele* Forel, 1904, *T. madeirense* Forel, 1895, *T. nigerrimum* (in the old large definition) and *T. simrothi* Krausse, 1911 (Berville et al 2013). We wanted to see if hydrocarbons can also be used in species discrimination, particularly in the *T. nigerrimum* group and we replaced it in the genus *Tapinoma* and some Dolichoderinae species from 11 countries: France, Germany, Switzerland, Belgium, Portugal, Spain, North Africa (Morocco, Algeria and Tunisia), Greece, and Italy.

## Methods

### Chemical analysis

Ten workers from each of the studied colonies were collected and killed by freezing. All the ants were immersed in 1 ml of hexane for 60 minutes, after which the ants were retrieved from the vials and the solvent evaporated. The samples were kept frozen at - 20°C until chemical analyses. For chemical analyses performed via a GC/MS-TQ Agilent (GC 7890B, MS 7000C, Agilent Technologies, Santa Clara, CA, USA), the samples were re-dissolved in 50 μl of hexane. Two µL of each extract were injected with an autosampler (Gerstel, Mühleim an der Ruhr, Germany) into an injector heated at 280 ºC in splitless mode and then in a column compound of 5% Phenyl - 95% Dimethylpolysiloxane (Zebron ZH-5HT inferno, 30 m × 0.25 mm × 0.25 µm, Phenomenex, Torrance, CA, USA). The gas vector was helium at 1.2 ml min-1. The temperature program was 2 min at 150°C, and then increasing at 5°C/min to 320°C, and 5 min hold at 320°C (total 41 min). The transfer line was set at 320 ºC. We used Electron Ionization source at 230 ºC with electron energy of -70 eV and a scan range of 40 – 600 m/z with 3.7 scans/s. Compounds were identified by their fragmentation pattern, compared to standard alkanes, library data, and Kovats retention indices. All compounds were included in the analyses. When it was not possible to estimate the amount of each co-eluted compound, they were treated as a single compound. Sterols and other contaminants like phthalates were not included.

All the % of HCs are provided as mean ± SE (Standard Error) in Table1. The data were analyzed using cluster analysis on % with Euclidean distances and Ward method (Statistica 8.0 program). We also calculated the equivalent chain length which indicates the mean of hydrocarbons length ECL = (Σ(%CnxXn))/100) where Cn is the number of carbons and Xn the % of this category. Martin et al 2019 called it Mean chain−length. ECL is not sufficient to discriminate species but is a good indication to classify them in different groups according to the length of hydrocarbons.

We did not analyze here hydrocarbons under C21 to avoid possible volatile compounds from the glands.

**Table1, Map (see suppl data). List of species and samples**, a total of 4 genus and 13 species from 11 countries (513 samples from 299 sites, from sea level to 2 600m in Sierra Nevada). Columns: Genus, species, country, Department, City, Date of collect, latitude, longitude(decimal World Geodetic System WGS 84), altitude, collectors and determinators, number of samples, reference if already known (Seifert et al 2017, Berville et al 2013, Gouraud & Kaufmann 2022).

- ***Tapinoma***: *T. madeirense* (n=27), *T. simrothi* (n=49), *T. erraticum* (n=76), T. *melanocephalum* (Fabricius, 1793) (invasive tropical from greenhouses, n=6), *T. pygmaeum* (Dufour, 1857) (n=11), *T. nigerrimum* group with the 4 species: *T. darioi* (n=23, including samples from Italy, the country of the type), *T. magnum* (n=193), *T. ibericum* (n=37), and *T. nigerrimum s.str*. (n=26). In this group, a group appeared separated from the others in Spain Mountains, supporting the presence of a possible new species, waiting for morphological and genetic analyses to be formally described (Seifert com. pers.). It was provisionally named *Tapinoma sp. Spain* (n=34). Unfortunately, we were not able to find *T. subboreale* Seifert, 2012 from France.
- *Dolichoderus quadripunctatus* (Linnaeus, 1771) (n=11).
- The Argentine ant *Linepithema humile* (Mayr, 1868) (n=12).
- *Bothriomyrmex corsicus* Santschi, 1923, a parasite of *Tapinoma* (n=8).

## Results and discussion

### 1. Hydrocarbons of the different species

Hydrocarbon profiles were all typical with carbons chains from C23 to C39 (see Suppl data Table2). We did not analyze hydrocarbons under C20 which are partially volatiles and not important in colonial recognition. There were mainly linear alkanes, di, and trimethyl alkanes. We found very few alkenes (<1%) except in *Bothriomyrmex* (75±19%). Alcohols and other substances were also rare at these extraction temperatures. We found a total of 174 different hydrocarbons across the species studied with 25 substances having more than 1% of the total hydrocarbons. Guillem (et al 2016) found 222 HCs across 12 *Myrmica* species. We verified that the hydrocarbon profiles presented by L. Berville et al (2013) correspond to our results for *T. erraticum, T. madeirense* and *T. simrothi*. It appeared that *T. nigerrimum* in their analyses was the recently redescribed species *T. magnum*.

Three distinct clusters appear corresponding to ECL <=27, ECL = 29-30, and one intermediate groups with ECL=27-34 (Fig. 1). The maximum ECL are for *Linepithema humile* (Lh ECL=34.26±0.53) and *Bothriomyrmex corsicus* (Bothrio, ECL=32.62±0.32). These are discussed below.

**Fig1.**
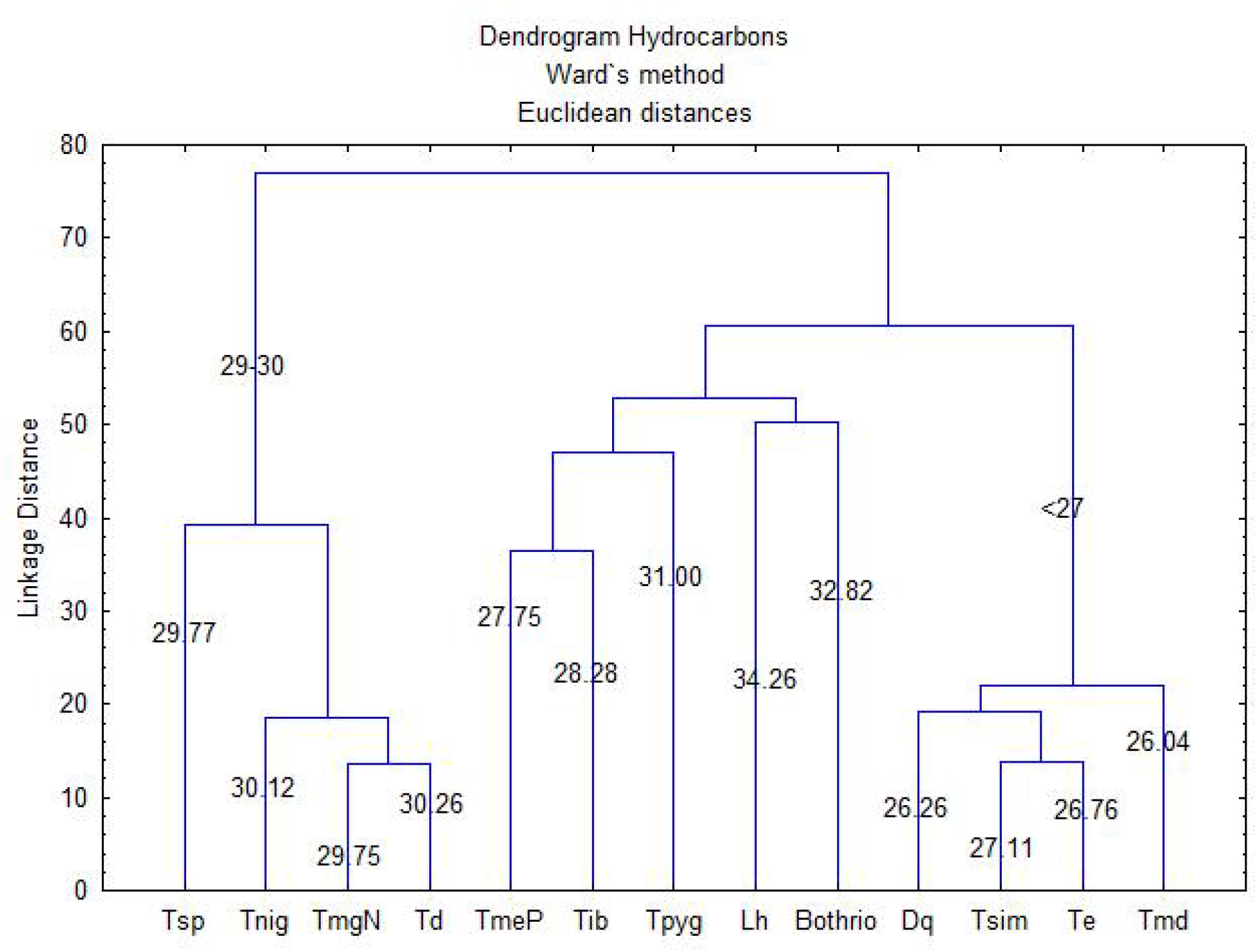
Dendrogram with Euclidean distances and Ward method on HCs % for all Dolichoderinae species from left to right: Tsp *T. sp. Spain*, Tn *T. nigerrimum*, TmN *T. magnum* natives, Td *T. darioi*, TmeP *T. melanocephalum* Paris, Tib *T. ibericum*, Tpyg *T. pygmaeum*, Lh *Linepithema humile*, Bothr *Bothriomyrmex corsicus*, Dq *Dolichoderus quadripunctatus*, Tsim *T. simrothi*, Te *T. erraticum*, Tmd *T. madeirense*. ECL are indicated on the figure.

### The first group (ECL <=27) consists of

*Dolichoderus quadripunctatus, Tapinoma erraticum, T. madeirense, T. simrothi* (see Fig2).

**Fig2.**
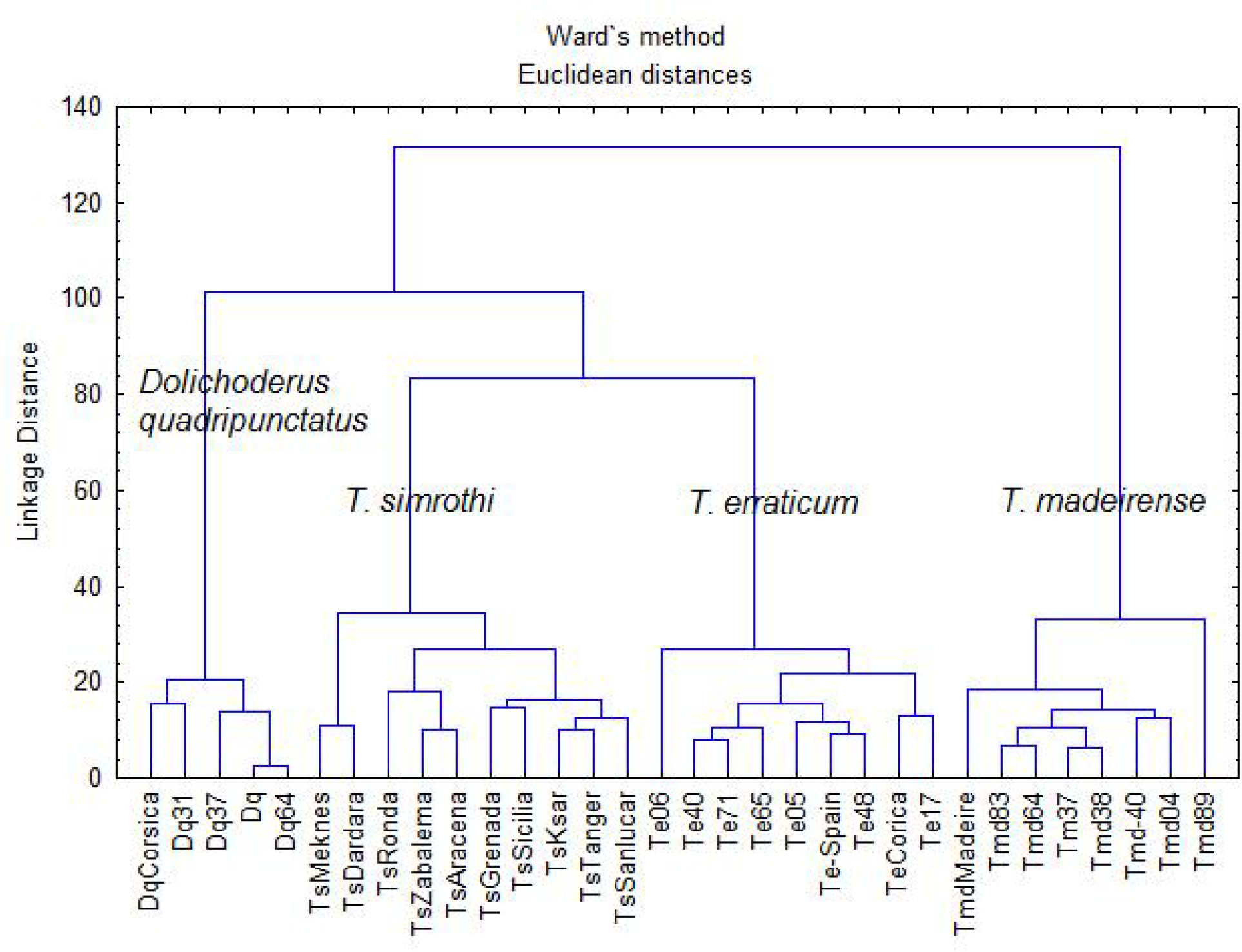
Dendrogram with Euclidian distances and Ward method on % for Dq *Dolichoderus quadripuncatus*, Ts *T. simrothi*, Te *T. erraticum, and* Tmd *T. madeirense*. Numbers indicate the department number for France, for example Dq37 id *D. quadripunctatus* Indre-et-Loire, and Corsica.

The 4 species appear to be clearly separated in Fig2.

#### Dolichoderus quadripunctatus

is the only arboricolous species. It is frequent everywhere in Europe, and present in 60 departments (and probably more) in France (Antarea, accessed on 10 Feb 2022, Blatrix et al 2013). It has a low ECL (26.26±0.13, n=11).

#### Tapinoma ibericum

is very frequent in South Spain (<41°) according to Seifert et al (2017): all Andalusia, found also in Portugal and in two places in Corsica (in red Fig3). ECL is 28.28±0.14. We did not observe differences between Spain, Portugal, and Corsica. It appears to become invasive in France, in a market gardening place near Pau (Meillon, 64), near Bordeaux (Saint-Médard-en-Jalles) and Lyon (Saint-Bonnet de Mure, B. Kaufmann leg). It is rare near Montpellier (1 site only at Mèze) (Centanni et al 2022). In Pozzuelo de Calatrava (Spain) where the *T. ibericum* holotypes were described by Santschi (Seifert et al 2017), only *T. magnum* was found (Ruano and Tinaut, leg). The two species are probably present in the same locality.

In Fig3 we analyzed T. ibericum and T. simrothi which are difficult to distinguish morphologically. They appear to be well separated based on CHs profiles.

**Fig3.**
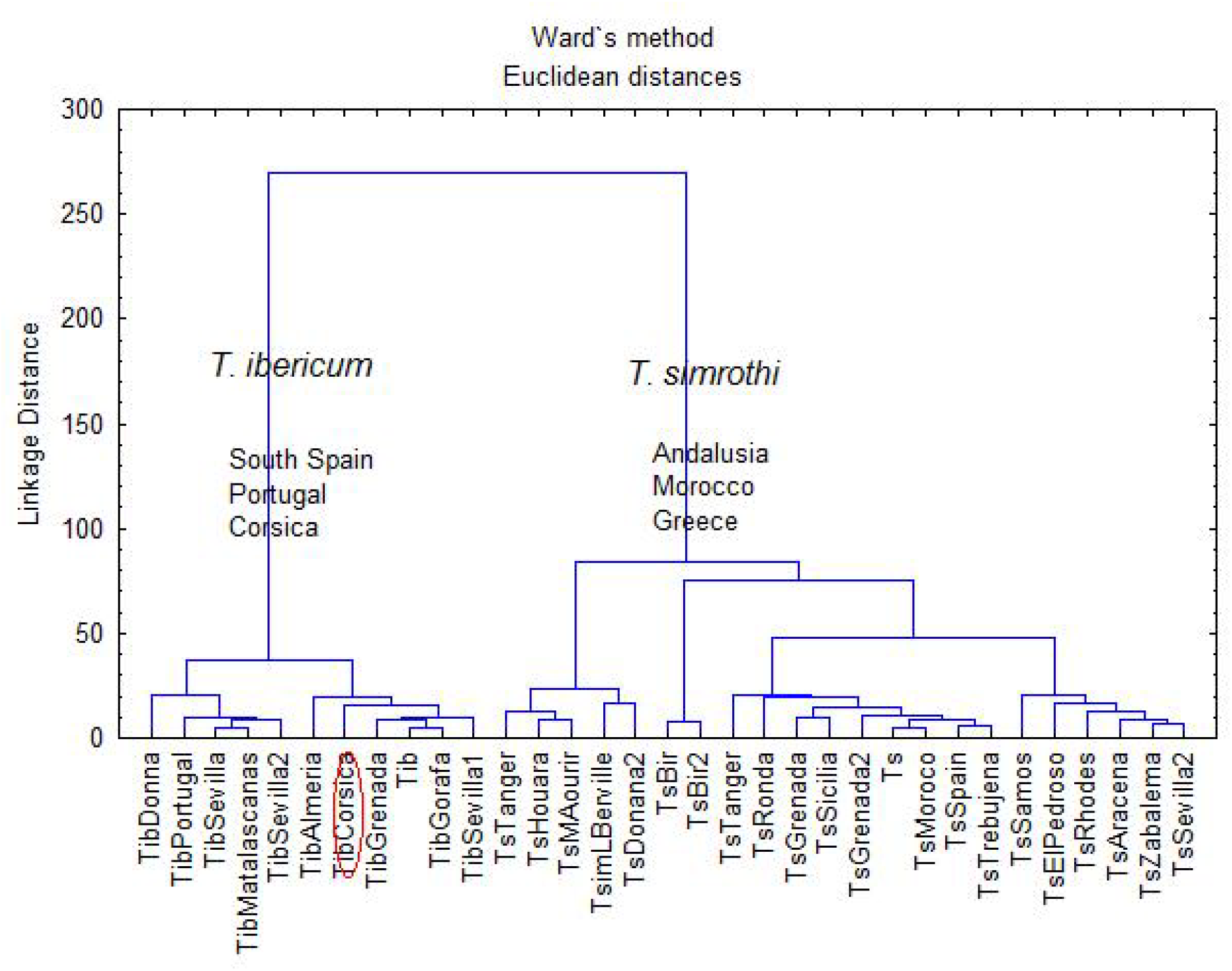
Dendrogram with Euclidean distances and Ward method on % for Tib *T. ibericum* (TibCorsica in Corsica in red) and Ts *T. simrothi*. Numbers indicate the department number for France

*T. ibericum* has to be now considered as an invasive species in France. It has the same HCs profile as natives ones. It will probably be found in many other places.

#### Tapinoma simrothi

is very frequent in Morocco under 500m (with one exception at 2 125m in Tichka col), frequent in Andalusia, in Greece (Salata & Borowiec 2018), and Sicilia. It has been found also in Corsica in two places (Antarea, accessed on 10 Feb 2022). According to Bernard (1979 and 1983), it proliferates in plantations in North Africa where it was probably introduced from Palestinia around 1890 since Forel did not find it in Algeria in 1869. ECL is low (27.11±0.57). The hydrocarbon profiles of this species are more heterogeneous than in *T. ibericum* (Fig3). This heterogeneity could reflect geographical structuring, which would be a good indication of the existence of cryptic species. In Lebanon for example, *T. simrothi phoenicium* is considered a subspecies of *T. simrothi* (Chanine-Hanna 1981).

#### Tapinoma erraticum and Tapinoma madeirense

In Fig4 we analyzed the two species T. erraticum and T. madeirense

There is a very good separation between *T. erraticum* and *T. madeirense* as found by Seifert (2012 – morphology and genetics) and Berville et al (2013 – morphology and HCs), although the two species have very similar ECLs: 26.04 ± 0.08 for *T. madeirense* and 26.76± 0.06 for *T. erraticum* (t test, *P=*0.30, NS). Surprisingly, the two species co-exist in some places like in Bléré (Fr: Te37 – Tm37 in red Fig4), which is a calcar dry place (both species confirmed by Xavier Espadaler pers.com.).

**Fig4.**
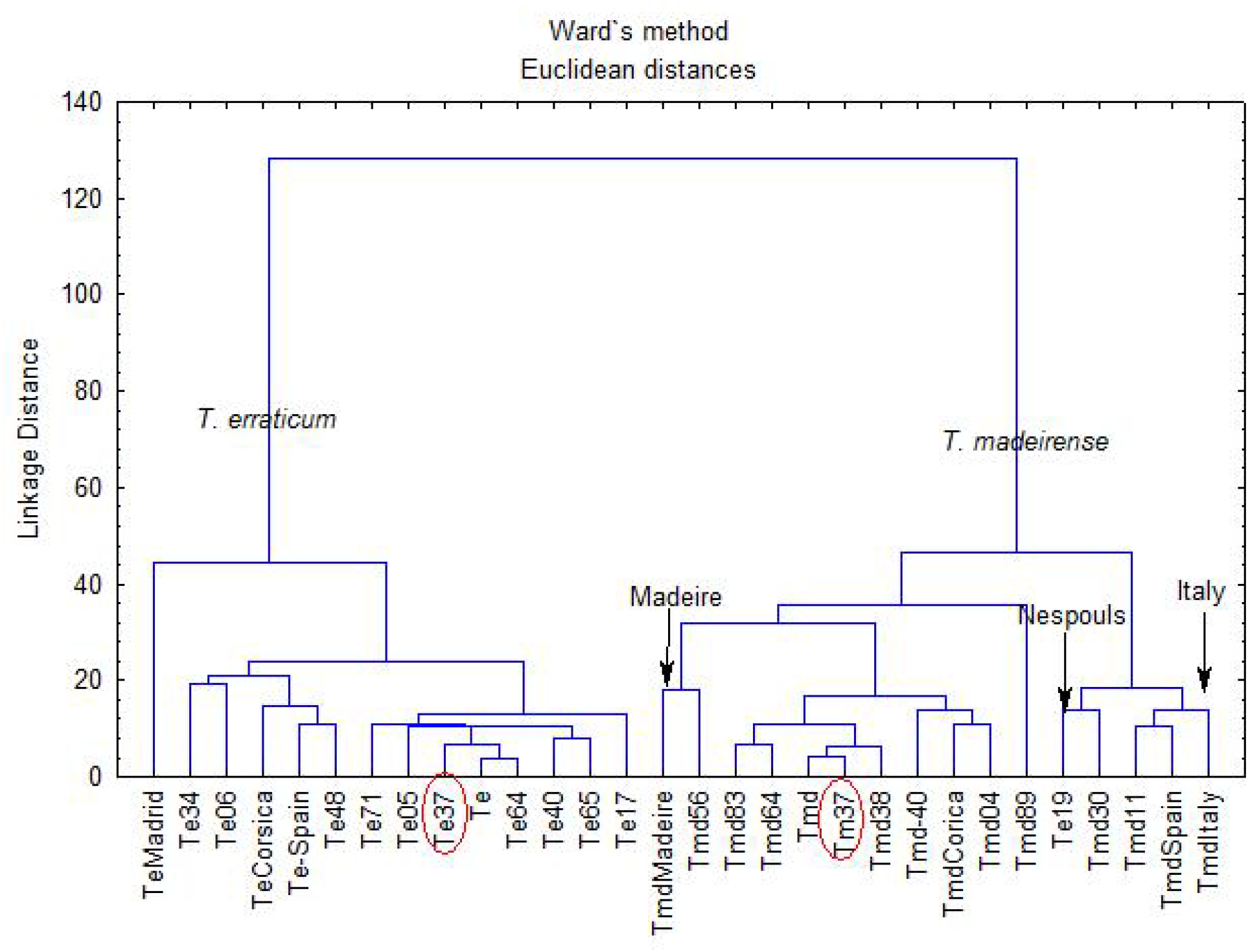
Dendrogram with Euclidean distances and Ward method on % for Te *T. erraticum* and Tmd *T. madeirense*. Numbers indicate the department number for France. Te37. *T. erraticum* and Tm37 *T. madeirense* from Blèrè (37) in red.

**Fig4b.**
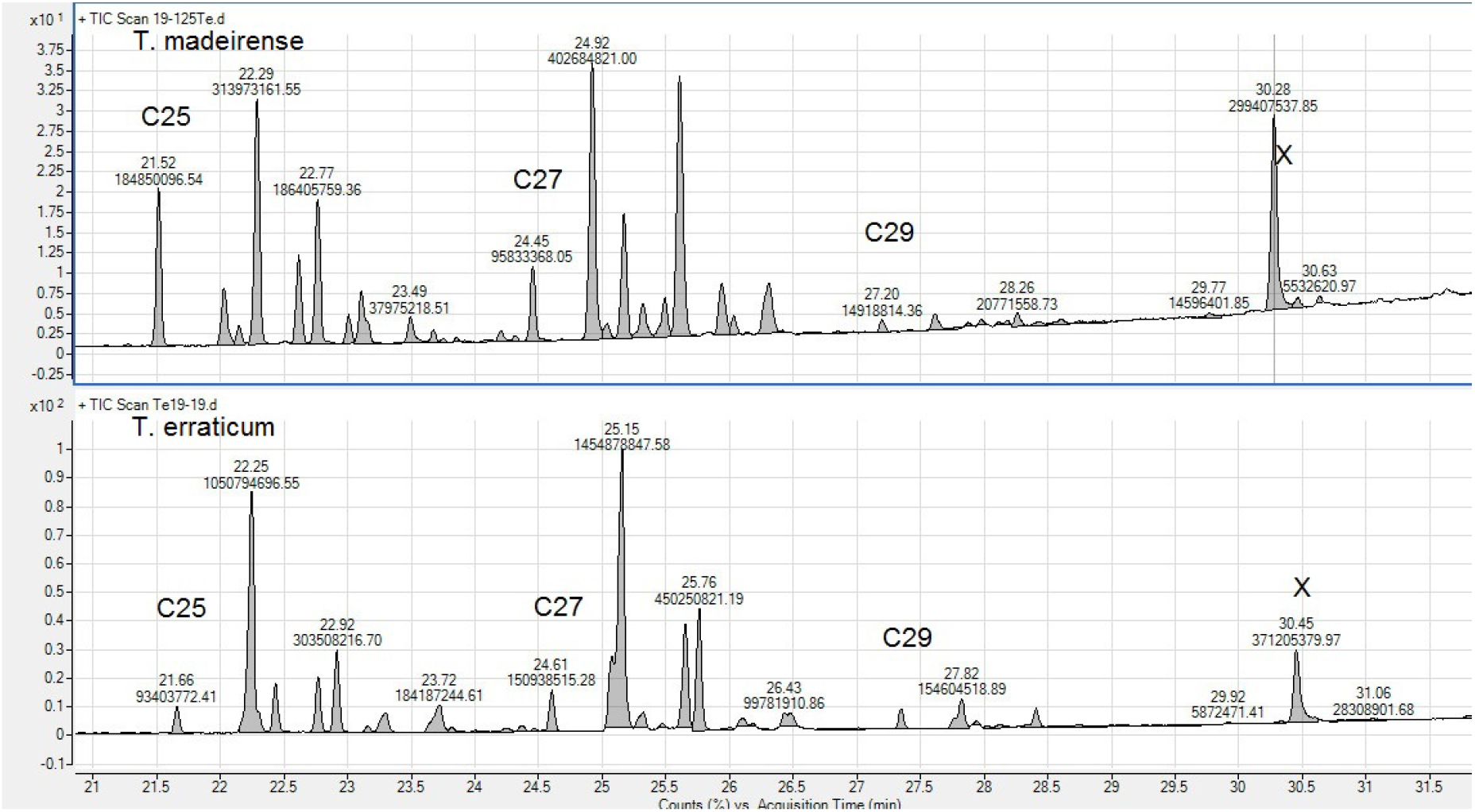
shows hydrocarbons profiles of the two species *T. erraticum and T. madeirense*, indicating very different ones (x=sterol)

We tried to collect neotypes of *T. erraticum* (Latreille, 1798) according Seifert (2012) in Nespouls, near Brive (19). In fact, they were *T. madeirense* (see Te19 within Tmad in Fig4). Probably the two species cohabit also in this place.

##### T. erraticum

was found in all departments surveyed in France (05, 06, 17, 34, 37, 40, 48, 56, 64, 65, 71, 20-Corsica), Madrid, and North Spain. This confirms its wide distribution across 85 departments in France (Antarea, accessed on 10 Feb 2022, see also Blatrix et al 2013). In the French Pyrénées mountains, it can be found up to 1 670m in the Gavarnie circus (65) and 2639m in Eyne (66, Lebas 2021) and at 2 100m according to Bernard (1986, p. 100), and up to 1 470m in Spain (Te-SP). In the Alps, it is signaled until 1 900m (Bernard 1983, p. 100). We found it at 1 400m (05 - Réallon). *T. erraticum* has been signaled in Algeria, Egypt, and Israel, but these could be misidentifications (Berville et al 2013). *T. erraticum* appears to extend in the Balkans, with two new sp. (Wagner et al 2018). It was found in Turkey but it could be a cryptic species, more samples are necessary to conclude (Kiran and Karaman 2020).

##### T. madeirense

was described from Portugal (Madeira island), which we confirmed (TmdMad). It is less frequent in France and mainly in the south (04, 11, 30, 40, 56, 64, 37, 38, 83, 89, 20-Corsica, 20 departments in South according to Antarea, accessed on 10 Feb 2022, see also Blatrix et al 2013), North Spain and Italy. In the North of France, it is not found in localities north of Yonne (89) and Indre-et-Loire (37). This species is not found above 900m.

In Fig5b it is interesting to see that the profiles are very identical, and therefore the species determination needs precise analysis. Nevertheless some differences appear clearly, for example at 38.10 min it is 8,10+8,14+8,16C30 (8,xC30 on the figure) for 3 species when it changes to 10,12+10,14C30 (10,xC30 on the figure for *T. sp. Spain T. nigerrimum* and *T. darioi* have also more hydrocarbons after C31, particularly 5,13+5,15+5,17C31 (5,xC31 on the figure). 8,x,xC32 on the figure is not representative due to the variability of the samples.

### The second group of species, with ECL C29 dominant

The second group of species, which is ECL C29 dominant (ECL 29-30), consists of 3 of the 4 known representatives of the *Tapinoma nigerrimum* group (*T. magnum TmgN* natives ECL is 29.74 ± 0.04 - we did not place here invasive ones, *T. darioi* ECL 30.26 ± 0.04, and *T. nigerrimum s.st*. ECL 30.12 ± 0.05.), and one new species (*T. sp. Spain* 29.83 ± 0.05), indicating that it is a complex of very close species. Surprisingly *T. ibericum* which was included in *T. nigerrimum* group by Seifert et al (2017) using morphometry and genetic data fall outside of this group as indicated before.

The 4 species were clearly separated according their hydrocarbons profiles (Fig5) but they do not differ according their ECL which are very close (Kruskall-Wallis P=0.000). D’Eustachio et al. (2019) analyzed alkaloids and volatiles ketones of *T. magnum and T. darioi* and confirmed the chemical difference between the two species. We did not analyze volatiles.

**Fig5.**
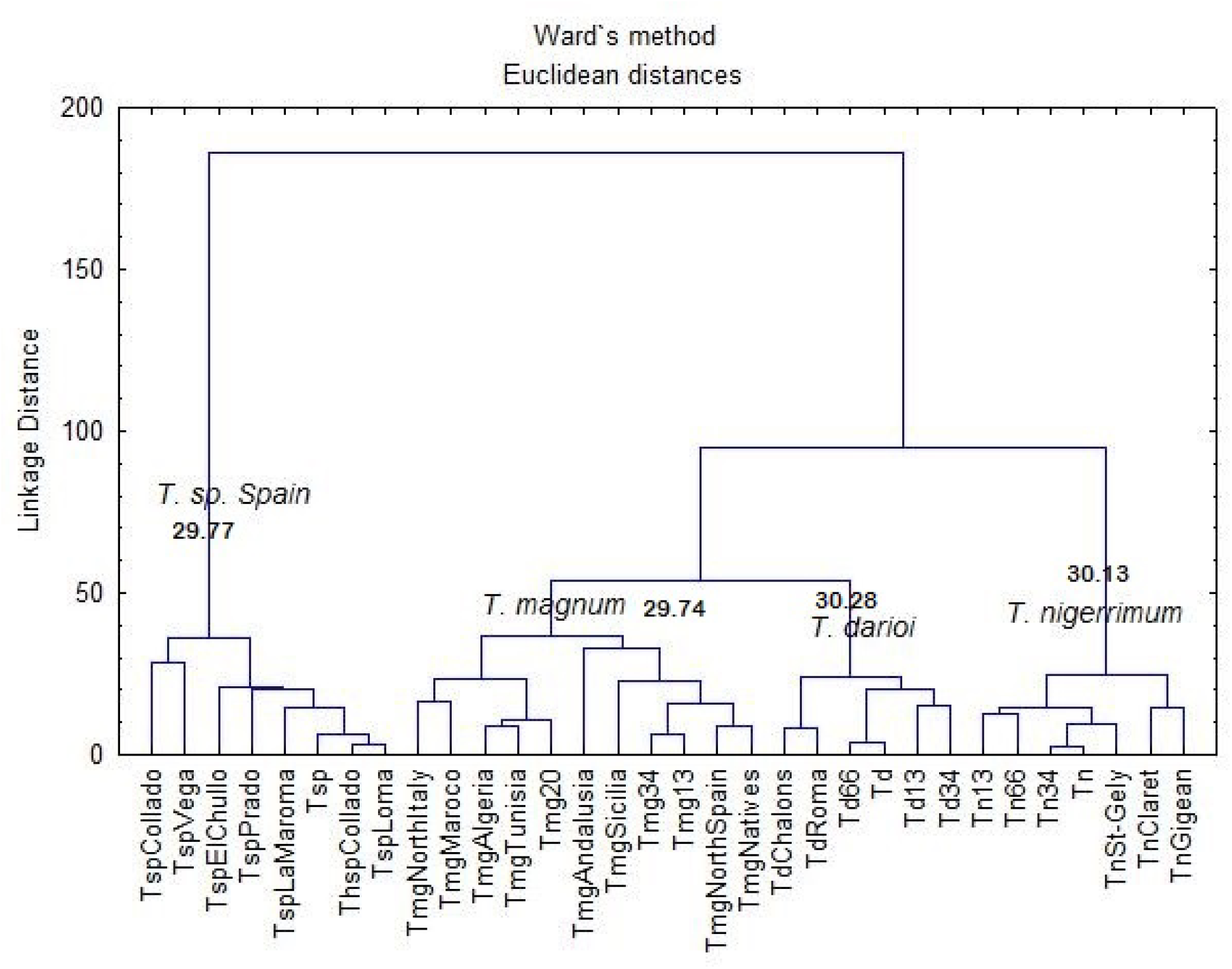
Dendrogram with Euclidean distances and Ward method on % for *Tapinoma* species of the *nigerrimum* group (Tsp *T. sp. Spain*, Tm *T. magnum*, Td *T. darioi*, Tn *T. nigerrimum*.

**Fig5b.**
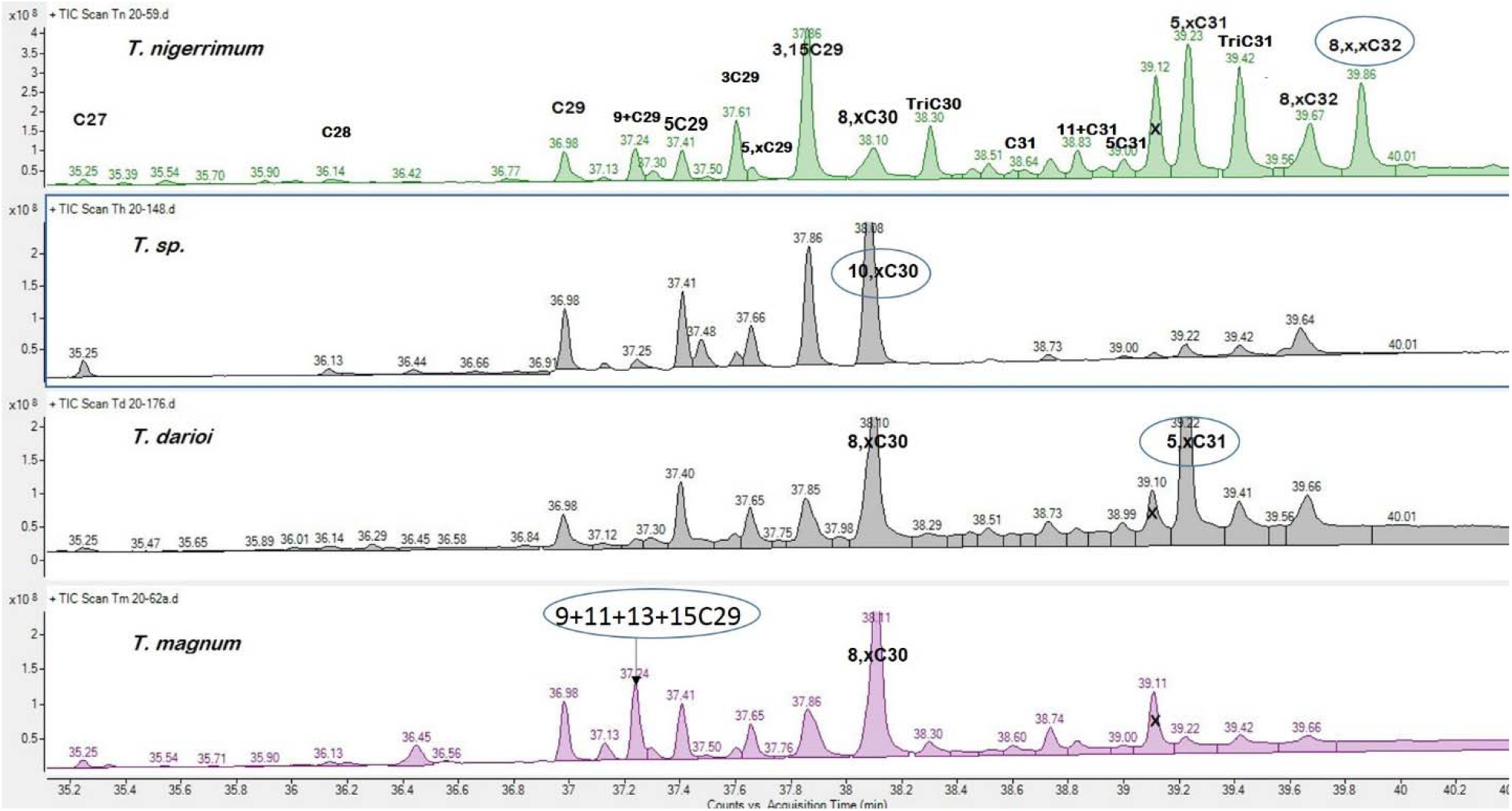
Chromatograms of the *T. nigerrimum* group. x is a sterol.

#### Tapinoma sp. Spain

This separated cluster, signaling a possible new species, is mainly from mountains in Sierra Nevada (>2 000m asl). ECL is 29.77 ± 0.04. It was also found in one locality in the mountains North of Madrid at Vega (Castilla) (980m, ECL = 29.42, n=2), but only one point, so it needs to be verified. It is chemically very different from the other species of *T. nigerrimum* group (Fig5c). It was considered previously as *T. nigerrimum* and therefore may be one more species to be added to the 72 endemic species for Spain (Tinaut & Ruano 2021). This needs morphometric and genetic analysis.

**Fig5c.**
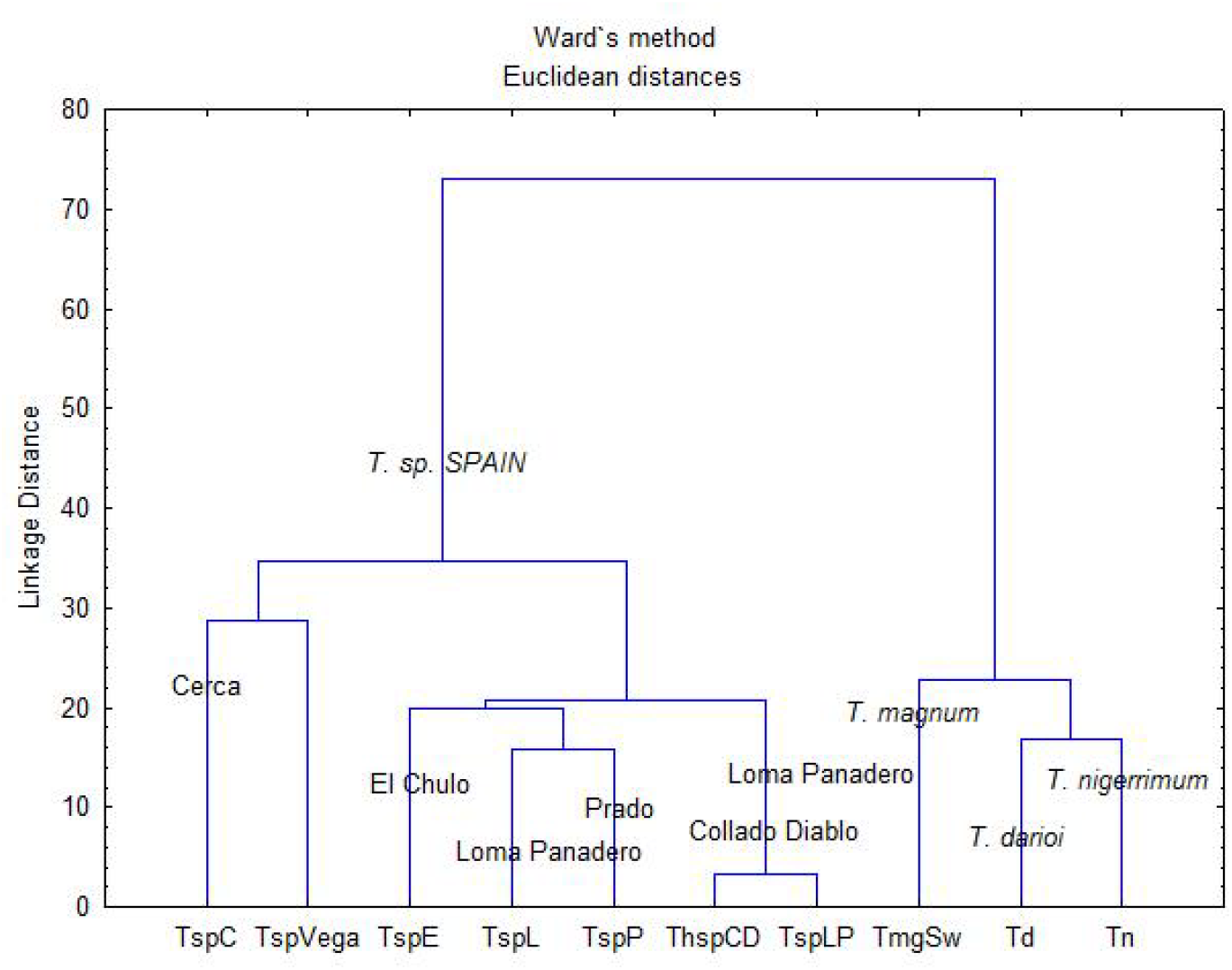

**Fig6.**
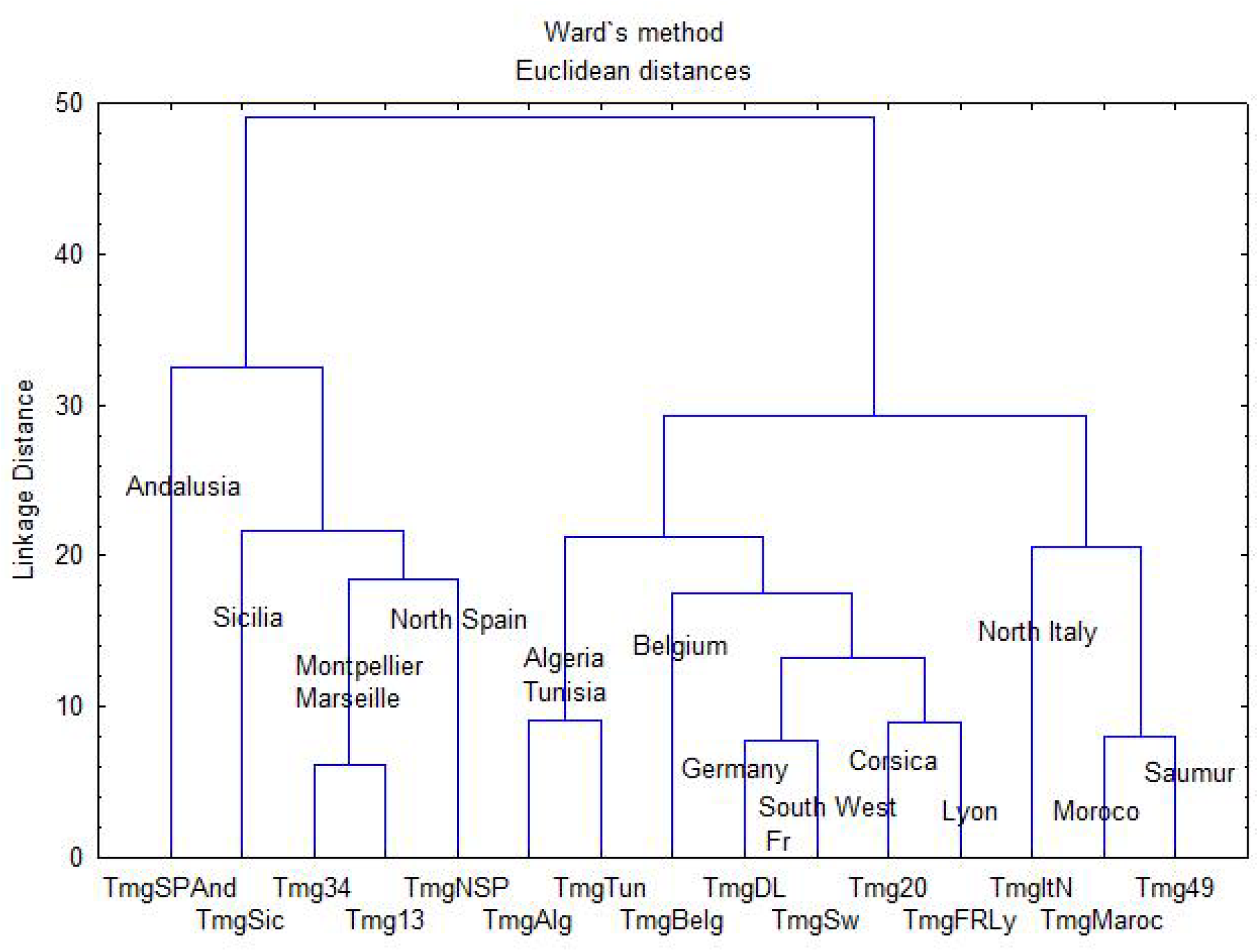
Dendrogram with Euclidean distances and Ward method on % for *Tapinoma magnum* of the different localizations.

##### Tapinoma darioi

is found in France in the Pyrénées-Orientales (66), Hérault (34), Aude (11), Marseille (13), Var (84), and in Italy in Roma (type locality, see Seifert 2012). *T. darioi* and *T. magnum* are occasionally found in the same localities in their invasive ranges. In Montpellier, *T. darioi* is frequent: 78 sites (8.42% of the studied sites) (Centanni et al 2022). It has been recently found in the Loire valley at Saint-Mars-du-Desert (44 - Gouraud & Kaufmann 2022).

##### Tapinoma nigerrimum *s.str*

is found in Europe on the Mediterranean coast from Provence to the Pyrénées-Orientales. It is frequent near the sea but is more generally found in lands up to 350m above sea level. Localities in Prades-Le-Lèzan and Gigean from Seifert are confirmed for this species, based on CHs. In Montpellier, *T. nigerrimum* is frequent: 197 sites (21.17% of the studied sites) and mainly observed on limestone plateaus and hills mostly covered with Mediterranean forests (Centanni et al 2022). It is also found in the mountains in North Madrid (800-1200m) and Italy (Genova).

#### Tapinoma magnum

In many papers, the ants called *T. nigerrimum* were probably *T. magnum*, for example in Fréjus (83-Fr) where colonies had up to 350 queens and 100% of the nests in the Piémanson beach (13-Fr) (Bernard 1983, p.100), which is not the case for the real *T. nigerrimum*. This was confirmed by Seifert (et al 2017) who found for example *T. magnum* in Fréjus beach.

*T. magnum* is now an invasive species spreading in many places in Europe and particularly in France like Britany (Gouraud & Kaufmann 2022, Lenoir et al 2022a). It has been found also in some places like a cemetery in Slovenia, it also probably arrived with plants (Bracko 2019).

- On the coast everywhere from Six-Fours (83), Cap d’Ail (06), Marseille, Saintes-Marie-de-la-Mer, Fos (13), near Montpellier (34), Girona (Spain), never higher than 20m. The three localities of Seifert (Le Grau du Roi, 2 spots in Saintes-Maries-de-la-Mer) are confirmed.
- Spain in Madrid region (700 to 1350m) and Andalusia (Doñana National Park in sand dunes). Seifert et al (2017) considered that *T. magnum* is rare in Spain, but either the number of samples was not sufficient, or it spreads rapidly.
- Corsica on the coast (3-4m) and higher in greenhouses (380-800m).*T. magnum* is becoming a pest in some places like Corsica for market gardening.
- Italy: Roma (57m) and Sicilia (900m).
- Morocco (more than 170m until 1 200m), Algeria (from sea level to 800m), and Tunisia (under 220m). *T. magnum* has been studied in the Algerian National Park where it represents 16% of all the ants (Labacci et al 2020).
- France: in Antarea it is found only 69 times in 13 departments (accessed 2 November 2022). In fact it now found in South West around Bordeaux (Galkowski 2008) and Arcachon (33), Dax (40), Agen (47), Sauvagnon and Arzacq-Arrizet near Pau (64), Bergerac (24), probably Toulouse (31).

It is invasive in the Loire valley, found by Gouraud and Kaufmann (2022): Saumur-49-where it is becoming a veritable pleague, Ancenis and Saint-Germain-sur-Moine, Ingrandes-Le-Fresne-sur-Loire; in the department 44: Le Croisic, Saint-Mars-du-Désert, La Suze-sur-Sarthe, Saint Nazaire, Batz-sur-Mer and Saint-Lyphard. It is also found in Lyon and Ternand (69), Bourg-en-Bresse (01), Molières (82) (Lenoir et al 2022a).

- Belgium: Ostende (Dekoninck et al 2015).
- Switzerland many places around Lausanne (Freitag and Cherix 2017).
- Germany: Edersheim, Ginsheim and Ingelheim (Seifert et al 2017).

Our data confirm Seifert et al (2012, 2017) results for *T. nigerrimum* group: *T. nigerrimum* is mainly more distant than 4 km from the coast but can be found near the sea (14% of the places). *T. darioi* is more present near the sea (80% - Siefert et al 2017). *T. magnum* is very present in degraded areas on human influence which is typical of invasive species. In Montpellier, *T. magnum* is not frequent: 6 sites (0.65% of the studied sites) and it is replaced by *T. darioi* (Centanni et al 2022).

We did not observe chemical differences between native and invasive colonies of *T. magnum*; the profiles are identical (Fig 7). This indicates that no dramatic changes in odour occur with migration. It was verified in colonies maintained in the laboratory for one or two years which kept their chemical profile contrarily to many other invasive species (Lenoir et al 2022b). Two groups appear within *T. magnum* which may correspond to two different genetic groups or different origins but the Euclidian distance is low (=50). This deserves further study.

**Fig7.**
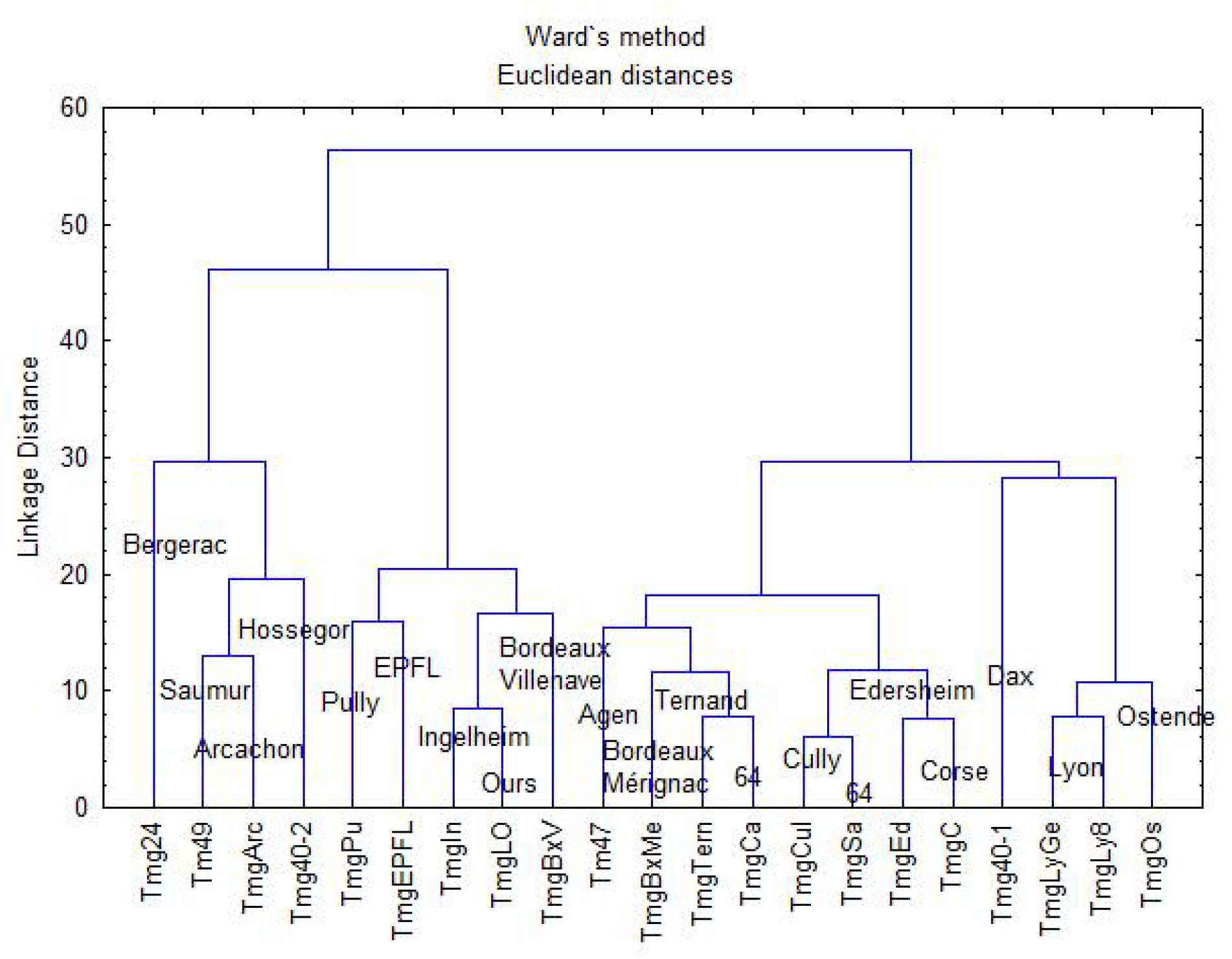
Dendrogram with Euclidean distances and Ward method on % for imported

In Fig7 we have the imported colonies plus Corsica. They present also two groups. We do not know the origins which can be different. For example, in Bordeaux there are possibly two origins as the profiles differ between Villenave and Mérignac.

We also tested the aggressiveness between colonies of *T. magnum* from different localities to see if this species form a unique giant colony like the Argentine ant. 10 ants were placed in a Petri dish and after 10 minutes we introduced one marked ant from another colony. The reaction of the ants were observed during 10 minutes, but generally very rapidly the result was obtained, either the introduced ant is accepted and exchanges with the other ones, or it is rejected and aggressed. In the case it was retrieved to prevent its death. The % of adoptions in aggression tests between two colonies were the following (n=10) : Sauvagnon / Lausanne 0% adoptions, Sauvagnon / Cully (Sw) 25% indifference and 75% rejections, Lausanne / Cully 100% adoptions, Lausanne / Bordeaux Mérignac 0%, Sauvagnon / Bordeaux Mérignac 0%, Sauvagnon first colony / new colony 100%, Sauvagnon / Caubios (5 km) 100%, Saumur zone A / zone B 100%, Arzaq zone1 (64) / Arzac zone 2 (200m) 100%, Arzacq / Sauvagnon 80% (same origin?).

The aggression between species is always maximum. For example we tested Sauvagnon / Meillon (*T. ibericum*) 0% adoptions, Sauvagnon / *Lasius niger* 0%. It indicates that T. magnum is very aggressive toward other species and explains probably why they exclude local species.

These results indicate that

- Colonies in a large cities like Saumur make a supercolony, and colonies from small distances like Lausanne and Cully are not aggressive, coming probably from the same plant importation.
- There is not a unique giant supercolony as aggression appears between various localities like Bordeaux and Lausanne, or Bordeaux and Sauvagnon.

**The third group (**ECL=27 to 34; Fig. 1) contains the four remaining species studied: *L. humile, T. melanocephalum, T. pygmaeum*, and *B. corsicus*.

### Linepithema humile

It is known as invasive species, found in South France in 13 departments (Antarea accessed 10 Feb 2022), but it seems to expand rapidly. It was found recently In Nantes city (Charrier et al., 2020). Hydrocarbons profiles of the argentine ant are well known, including the queens (see for example Blight et al 2012, Abril et al 2018, Buellesbach 2018 for California). Three supercolonies are known: Main European, Corsican and Catalonian according to Blight et al (2012). We analyzed ants of the Main European Supercolony from Italy and Spain. This species has the higher ECL (34.26 ± 0.53, n=12) of all studied Dolichoderinae ants with mainly C35 (12.56%±2.04), C36 (30.61%±3.44) and C37 (26.90%±4.07). These long-chain hydrocarbons which protect against desiccation may allow *L. humile* to support very dry climates. Long-chain compounds are generally thought to enhance desiccation resistance (reviewed for example by Gibbs 1998).

### Tapinoma melanocephalum

It is a very frequent tropical species (see the taxonomic position in Guerrero 2018) and one of the most invasive ant species in the world.

It is also an invasive species found in greenhouses of many tropical botanical gardens in Europe (Blatrix et al 2018). We found it in the Jardin des Plantes (Muséum d’Histoire Naturelle Paris) and in the botanical garden in Villers-lès-Nancy. It was also found in the University city of Villeurbanne (69, T. Klaftenberger, 16 August 2021), in Roubaix (Anaïs Tamelikecht / Agnès Villain, 10 November 2020), but in this last case, it needs to be verified. According to Antarea it has been found in 13 departments (accessed 10 Feb 2022). It has been signaled in a building in Liege (Dekoninck et al 2006) and in Czech Republic (Klimes and Okrouhlik 2015).

We studied ants from the Jardin des Plantes in Paris. ECL is 27.75±0.04 (n=6). They have a very simple profile with only 9 HCs >1% (C27 16.31%±1.50, 9+11+13C27 9.89±0.59, 3C27 24.61±1.86, C29 13.62±1.14, 9+11+13+15C29 21.18±1.35).

This species may be composed of several species as Siefert (2022) found a new species *T. pithecorum* in Indo-pacific region.

### Tapinoma pygmaeum

It is a rare *Tapinoma* species described from Saint-Sever (40, Landes, Emery 1912), rediscovered in France in 1999 (Péru 1999). It is found in 22 French departments (Antarea, accessed on 10 Feb 2022). We found it near Chartres and near Tours (La Riche and Montlouis). Hydrocarbons have long-chain molecules: ECL=31.00±0.30 (n=11).

### Bothriomyrmex corsicus

It is a rare parasite species found in places with a high density of *Tapinoma* in Pyrénées-Orientales and near Tours (Bléré). Antarea indicated it on 17 departments mainly in the South (accessed 10 Feb 2022). We found only pure *Bothriomyrmex* colonies. ECL is very high (32.62±0.32, n=8). It has a hydrocarbon profile very particular with a great quantity of alkenes (74.99±18.81%), in some individual ants it was more than 90% whereas other Dolichoderinae has only very few alkenes. This species has been studied only for volatile compounds. The *Bothriomyrmex* queens are able to enter the *Tapinoma* colony with a ketone produced only by the queen (Lyod et al 1986). The abundance of alkenes in workers may also be related to parasitism. The inquiline ant *Myrmica karavajevi*, parasite of *Myrmica scabrinodis*, used two adaptations to be admitted in the host colony, it smells the host queen odor but also produces sounds similar to the host ants (Casacci et al 2021). The total quantities of alkenes are more important in the *M. karavajevi* parasite queens (16.68%) compared to *M. scabrinodis* workers (8.58%) but they are very far from the *Bothriomyrmex* quantities.

## Conclusions - Discussion

Cuticular hydrocarbons of Dolichoderine ants are classical with carbons chains from C23 to C39. All species can be identified with their specific profile and a possible new species is identified. It is interesting to see that cuticular hydrocarbons profile is an efficient tool to determine Dolichoderine ant species and particularly in the *T. nigerrimum* group where morphology is very difficult and reserved to good specialists, and when genetic data are not possible. The parasite *Bothriomyrmex* is very different from all other species with a lot of alkenes, probably linked to the parasite life, but it needs to be discussed.

The 4 species of *T. nigerrimum* group described by Seifert (et al 2017) are well discriminated with hydrocarbons profiles. Surprisingly they were divided into two clearly separated groups: a first group with 3 species: *T. magnum, T. darioi, T. nigerrimum s.str*. and the new *T. sp Spain* and *T. ibericum* in another different group. *T. magnum and T. darioi* live in different places and form supercolonies (Cenatti et al 2022). It indicates that morphometric plus genetic analyzes versus hydrocarbons can classify the species differently. It is interesting to note that *T. magnum* forms very large supercolonies in cities, but not giant supercolonies like *Linepithema humile*.

*T. ibericum* and *T. simrothi* are well differentiated and have a large distribution in Spain and North Africa. *T. ibericum* is mainly from Spain while *T. simrothi* from Morocco (and Corsica).

*T. erraticum* and *T. madeirense* have a very large distribution. They can be present in the same habitat but probably have different microclimatic preferences. According to Claude Lebas (pers. comm.) *T. madeirense* lives only in deadwood.

*T. melanocephalum* is imported in France and found in most all green-houses and must be surveyed in city flats as it could become invasive.

*Tapinoma pygmaeum* is a rare species with a particular microhabitat, it is well separated from all other ones with HCs and morphology.

### Perspectives

More analyses are necessary to analyze relations between cuticular hydrocarbons composition and adaptations to climate. It is generally accepted that ants can plastically adjust their profile to acclimate to different conditions. Warm-acclimated individuals generally show longer n-alkanes and fewer dimethyl alkanes. Dry conditions result in more n-alkanes and fewer dimethyl alkanes for workers, probably due to better resistance to desiccation (Menzel et al 2017, 2018). *Aphaenogaster iberica* in the Sierra Nevada mountains show also differences in n-alkanes due to the elevation (Villalta et al 2020).

It will be interesting to follow the progression of some species, mainly *T. magnum* but also *T. darioi and T. ibericum. T. magnum* and *T. darioi* are native in the South east of France but in these regions they are becoming invasive for example in Montpellier region (Centanni et al 2022). Two hydrocarbons profiles of *T. magnum* appear, it will be interesting to see if they have genetic differences.

## Supporting information

Map of collected ants

## Acknowledgments

to people who collected and/or determined ants. In France: Sulpice Clément (Dep 01), Olivier Blight (OB) and Hélène Dumas (13), Mathieu Lenouvel (MLe, Laboratoire LMC Sarrolla-Carpopino, Corse, 20), Benoit Cailleret (BC, Association Areflex, San Giulano, Corse), Christian Foin (24), Henri Cagniant and Luc Passera (31), Rumsais Blatrix (RB, CNRS Montpellier, 34), Damien Alcade (47), Loïc Bidaud (Angers Municipality) and Christophe Meunier (Saint-Germain-sur-Moine, 49), Sylvie Elhorga, Denis Lafaille and Mathilde Lenoir (64), Claude Lebas (CL, 66), Bernard Kaufmann (BK, LEHNA Lyon, 69), Quentin Rome (Muséum d’Histoire Naturelle Paris) and Romain Péronnet (iEES Paris), Christian Champagne (82), Olivier Blight (OB, IMBE Avignon) and Hélène Dumas (84), Gérard Renaud (Monaco). In Spain: Xavier Espadaler (XE, Universidad Autónoma de Barcelona), Xim Cerdà and Elena Angulo (Estación Biológica de Doñana), Mariola Sivestre and Francisco M. Azcarate (Universidad Autonoma de Madrid), Francisca Ruano and Alberto Tinaut (Universidad de Granada). In Portugal Vera Zina (University of Lisboa). In Switzerland Cleo Bertelsmeier (CB) and Jérôme Gippet (JG) (Université de Lausanne). In Morocco Ahmed Taheri (AT, Faculté des Sciences de Tétouan). In Italy Alberto Fanfani (Universita di Roma). In Belgium Deconinck Wouters (Institut Royal des Sciences Naturelles de Belgique).

We particularly thank Christophe Galkowski (CG) who determined morphologically a lot of samples and collected *Tapinoma* in the region of Bordeaux. Bernard Kaufmann and Hugo Darras determined some samples using mitochondrial DNA. Special thanks to Laurent Keller who permitted to analyze samples collected in North Africa by Claude Lebas. Thanks to Francisca Ruano and Alberto Tinaut who discovered the possible new *Tapinoma* species and Bernhard Seifert who confirmed that it is a new species to be described later (*T. sp. Spain*).

## Participation

Ant collection: Alain Lenoir and all persons indicated in the acknowledgments. GC-MS utilization Elfie Perdereau and A. Lenoir. Analysis of data A. Lenoir and Laurence Berville. Redaction A. Lenoir, Elfie Perdereau and Laurence Berville. Thanks to Hugo Darras, Claude Lebas, Rumsais Blatrix, Francisca Ruano, Julia Centanni, Jean-Luc Mercier, Bernard Kaufmann, and Cleo Bertelsmeier for their helpful and critical comments on previous versions of the manuscript.

## Supplementary data

- Map: distribution of the ants collected
- Table 1: List of species and samples
- Table 2: Hydrocarbons composition of the species

## Websites

- Antarea: http://antarea.fr/fourmi/?

