## Supplementary figures and images for "Chemotaxonomy of *Tapinoma* and some Dolichoderinae ants from Europe and North Africa"

### Map of collected ants

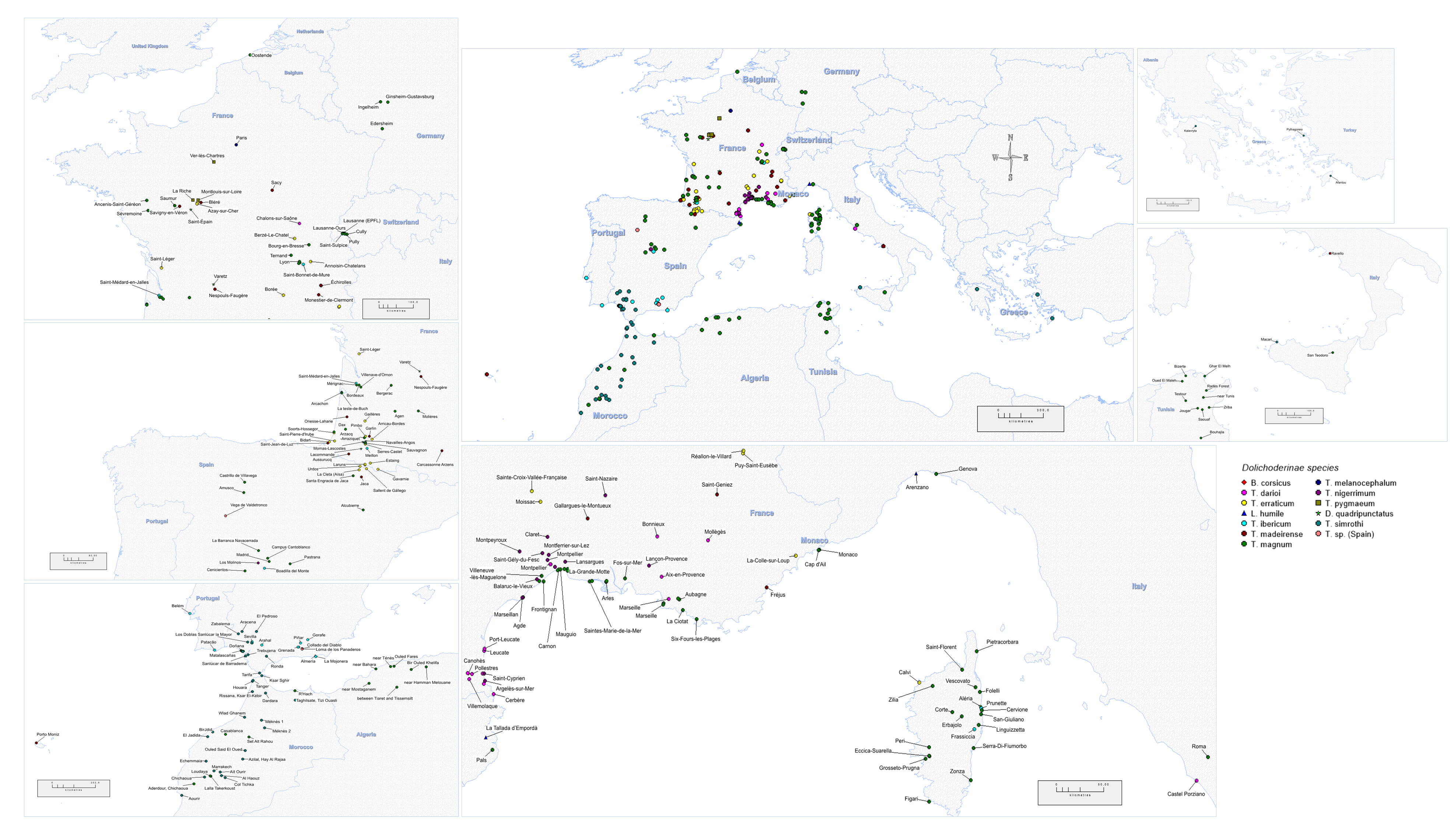
